# Glymphatic System Is Enhanced in Anti-NMDAR Encephalitis and Facilitates Antibody Influx and Clearance in the Brain

**DOI:** 10.64898/2026.10.15.759034

**Authors:** Dongmei Wang, Jingsi Liu, Fenfen He, Yueyang Liang, Yixin Guo, Yaqin Zhang, Donghua Chen, Zijian Xu, Yongming Wu, Kaibiao Xu

## Abstract

**Background:** Anti-N-methyl-D-aspartate receptor (anti-NMDAR) encephalitis is an antibody-mediated neuroinflammatory disease. How pathogenic autoantibodies enter and are cleared from the brain parenchyma remains poorly understood. The glymphatic system (GS) mediates cerebrospinal fluid (CSF)-interstitial fluid (ISF) exchange and waste removal, yet its role in anti-NMDAR encephalitis is undefined.

**Methods:** We used polypeptides GluN1_356-385_ induced active-immunization anti-NMDAR encephalitis mouse model. Glymphatic influx and efflux were evaluated via fluorescent tracer delivery combined with real-time imaging and ex-vivo quantification. Perivascular aquaporin-4 (AQP4) polarization, brain IgG deposition, NMDAR levels and blood-brain barrier (BBB) integrity were analyzed. GS function was pharmacologically modulated with AER-271 (an AQP4 inhibitor) or dexmedetomidine (Dex), and deep cervical lymphatic vessels (dCLV) were ligated in a passive-transfer model (PTM). Behavioral performance was characterized by multiple neurobehavioral tests.

**Results:** Both glymphatic influx and efflux were enhanced in anti-NMDAR encephalitis mice, accompanied by upregulated antibody entry into and clearance from the brain. However, the cerebral IgG deposition was increased. These changes were associated with increased perivascular AQP4 polarization, with no alterations in overall AQP4 abundance or BBB tight-junction integrity. AQP4 inhibition with AER-271 attenuated glymphatic influx and efflux, reduced hippocampus IgG deposition, and partially ameliorated behavioral deficits without restoring hippocampus NMDAR levels. Conversely, dCLV ligation in the PTM mice increased brain retention of patient-derived IgG and exacerbated anxiety-like behavior. Dex enhanced antibody clearance and reduced hippocampus IgG deposition but did not improve behavioral outcomes.

**Conclusion:** The GS is dynamically upregulated in anti-NMDAR encephalitis, driven by increased perivascular AQP4 polarization. The enhanced glymphatic influx facilitates antibody entry into the brain parenchyma and exacerbates neuronal injury, whereas enhanced efflux provides a compensatory route for antibody clearance. Rather than broadly enhancing glymphatic activity, precise tuning of this pathway may present a promising non-immunological therapeutic strategy for autoimmune encephalitis.

## INTRODUCTION

Anti-N-methyl-D-aspartate receptor (anti-NMDAR) encephalitis is a severe, antibody-mediated neuroinflammatory disorder characterized by autoantibodies against the GluN1 subunit of the NMDAR [1]. The pathogenesis is driven by the penetration of these pathogenic antibodies into the brain parenchyma, where they bind to and internalize NMDARs, leading to synaptic dysfunction and diverse neuropsychiatric manifestations [2]. Current therapeutic approaches mainly aim to eliminate circulating peripheral antibodies. The mechanisms governing the entry of antibodies from the cerebrospinal fluid (CSF) into the brain parenchyma and their subsequent clearance remain poorly understood. The traditional view assumes that antibody entry occurs via passive leakage across a disrupted blood-brain barrier (BBB) [3]. However, clinical observations suggest that approximately 70% of patients maintain an intact BBB, and 50%–65% exhibit normal cranial MRI findings without obvious BBB breakdown or vasogenic edema [4, 5]. These discrepancies suggest that BBB leakage alone is insufficient to explain antibody brain entry, highlighting the need to explore alternative, possibly active transport mechanisms.

The glymphatic system (GS) is a recently discovered brain-wide waste clearance network that facilitates the exchange of cerebrospinal fluid (CSF) and interstitial fluid (ISF) via the perivascular space (PVS) [6]. The GS is anatomically composed of three key components: the periarterial CSF influx pathway, the astrocyte-driven convective fluid transport dependent on polarized aquaporin-4 (AQP4) water channels on astrocytic endfeet, and the perivenous ISF efflux pathway that drains into the meningeal lymphatic vessels and deep cervical lymph vessels (dCLVs) [7, 8]. This system plays an important role in clearing metabolic waste products, including β-amyloid and tau proteins, as well as inflammatory mediators, thus maintaining central nervous system (CNS) homeostasis [9, 10].

GS dysfunction contributes to the pathology of many neurological conditions, including Alzheimer’s disease (AD), Parkinson’s disease (PD), stroke, and traumatic brain injury (TBI), where it leads to the accumulation of neurotoxic proteins and the exacerbation of neuroinflammation [11, 12]. In the CNS inflammation, recent studies have shown that the GS closely interacts with BBB properties. The BBB, composed of endothelial cells, tight junctions, and a basement membrane, forms the first barrier against circulating immune cells and macromolecules. Meanwhile, a second barrier, the “glia limitans”, formed by astrocytic endfeet and their basement membrane, is also a critical component of the BBB and is an integral part of the GS pathway [13, 14]. The crosstalk between the GS and BBB is therefore an area of interest in understanding neuroinflammatory disorders [15].

GS function relies on polarized AQP4 distribution on astrocytic endfeet and can be dynamically modulated by a variety of factors, including inflammation, sleep, and pharmacological agents such as dexmedetomidine (Dex)[16]. As an α2-adrenergic receptor agonist, Dex is known to enhance GS function. It increases CSF tracer influx and clearance in rodent models by lowering central noradrenergic tone and activating the PI3K/AKT/ΔFosB/AQP4 signaling pathway [17, 18]. Selective AQP4 inhibitors, TGN-020 and AER-271, can suppress GS function[19–21]. These pharmacological tools highlight the therapeutic potential of the GS, either by promoting or inhibiting its activity.

In anti-NMDAR encephalitis, preventing the entry of the antibody into the brain parenchyma and enhancing the clearance of the antibody might provide new treatment strategy. Given the GS’s role in fluid and matter transport, we hypothesized that it might play a dual role in the pathophysiology of anti-NMDAR encephalitis. The GS may act as a conduit that brings pathogenic antibodies into the brain parenchyma; on the other hand, it might also serve as an important route to clear these antibodies and inflammatory mediators. In this study, we took advantage of the well-validated mouse models of anti-NMDAR encephalitis to elucidate GS-related mechanisms and explore whether targeting this pathway could serve as a novel non-immunological strategy to modulate antibody entry and removal.

## METHODS

### Animals

Eight-week-old female C57BL/6J mice were purchased from Rise Mice Biotechnology Co., Ltd. (Zhaoqing, China). Animals were allowed one-week acclimatization before experimental procedures. Mice were housed under specific-pathogen-free (SPF) conditions at 22 ± 2 °C with 55 ± 10 % humidity in the Laboratory Animal Center of Nanfang Hospital, Southern Medical University. All mice were maintained on a 12h light-dark cycle with free access to food and water. All animal experiments were approved by the Animal Ethics Committee of Nanfang Hospital, Southern Medical University (approval No. IACUC-LAC-20240903-003). Experiments were performed following AAALAC standards and the ARRIVE 2.0 reporting guidelines.

### Active immunization

The active-immunization mouse model of anti-NMDAR encephalitis was generated as previously described with minor adjustments (**Fig. 1A**) [22]. Synthetic polypeptides GluN1_356–385_ (LQNRKLVQVGYNGTHVIPNDRKIIWPGGE) was dissolved in PBS to a concentration of 2 mg/ml and emulsified 1:1 with complete Freund’s adjuvant (CFA) containing 600 µg heat-killed *Mycobacterium tuberculosis* H37Ra (final concentration 8 mg/ml). Mice received subcutaneous injections at two dorsal sites, 200 µg polypeptides per mouse. Booster immunizations were administered at weeks 4 and 8. Pertussis toxin, 200 ng per mouse, was injected intraperitoneally immediately after each immunization and repeated 48 h later. Sham-control animals received PBS emulsified with adjuvant using the same protocol, instead of the polypeptides.

**Figure 1.**
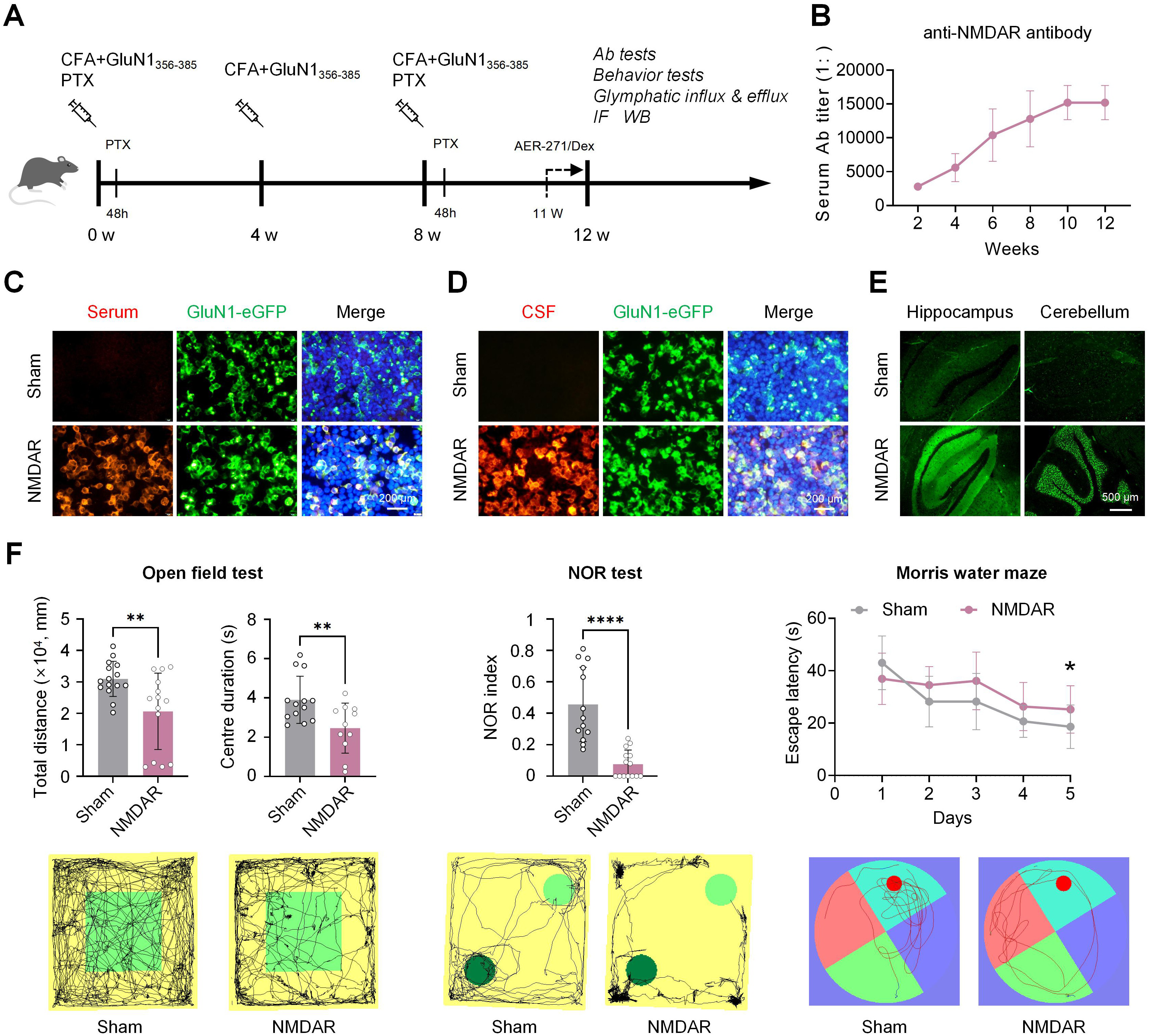
Establishment of the active-immunization mouse model of anti-NMDAR encephalitis. (A) Experimental workflow for generating and analyzing the GluN_356-385_ polypeptides-induced active-immunization anti-NMDAR encephalitis mouse model (NMDAR group). Mice receiving PBS instead of GluN_356-385_ served as Sham controls. When applicable, AER-271 or Dex treatment was administered every two days starting at week 11. (B) Longitudinal monitoring of serum anti-NMDAR antibody titers across experimental time points. (C, D) CBA assay for detection of anti-NMDAR antibodies in serum (C) and CSF (D). Anti-NMDAR antibody signal is shown in red, overexpressed GluN1-eGFP in green, and DAPI stained nuclei in blue. Scale bars, 200 μm. (E) TBA assay for detection of anti-NMDAR antibodies in serum (green). Scale bar, 500 μm. (F) Behavior assessments performed at week 12. In the open-field test, anti-NMDAR encephalitis mice exhibited reduced total travel distance (Sham, n = 15; NMDAR, n = 14) and shorter time spent in the central zone (Sham, n = 13; NMDAR, n = 11). The novel object recognition index was lower in anti-NMDAR encephalitis mice (Sham, n = 13; NMDAR, n = 14). Escape latency in the Morris water maze was prolonged in anti-NMDAR encephalitis mice (Sham, n = 13; NMDAR, n = 14). Representative movement-tracking plots are shown in lower panels. CBA, cell-based assay; CFA, complete Freund’s adjuvant; CSF, cerebrospinal fluid; Dex, dexmedetomidine; NMDAR, N-methyl-D-aspartate receptor; NOR, novel object recognition; PTX, pertussis toxin; TBA, tissue-based assay. \**P* < 0.05, \*\**P* < 0.01, and \*\*\*\**P* < 0.0001.

### Detection of anti-NMDAR antibodies by cell-based assay (CBA) and tissue-based assay (TBA)

Serum and CSF were collected from immunized mice. For CBA assay, HEK293T cells transiently transfected with pcDNA3.1-GluN1-eGFP plasmids were fixed with 4% paraformaldehyde (PFA) in PBS for 10 min, permeabilized with 0.3% Triton X-100 in PBS for 10 min, and blocked with 2% BSA in PBS for 30 min, at room temperature (RT). Cells were incubated with serial-diluted samples for 1 h at RT. After washing with PBST, cells were incubated with Alexa Fluor 568-conjugated goat anti-mouse IgG antibody (A11004, ThermoFisher, USA; 1:1,000) for 1 h at RT. Images were taken by using an inverted fluorescence microscope (IX73, Olympus, Japan).

TBA was performed to verify antibody binding toward native brain antigen. Frozen brain sections (8 µm) from naïve mice were fixed in ice-cold acetone for 10 min and blocked as described above. Sections were incubated overnight with mouse serum diluted 1:50 at 4 °C. After washes and secondary antibody staining, the sections were imaged.

### Primary neuron immunostaining

Primary rat hippocampal neuronal cultures were conducted as described previously[23]. To test antibody reactivity against primary neurons, mouse serum or CSF samples from patients with anti-NMDAR encephalitis were applied to the cultured neurons at 10-14 days in vitro (DIV) for 24 h. After fixation with 4% PFA in PBS for 10 min at RT, cells were subjected to immunostaining and imaged as CBA. The neurons were costained with anti-microtubule-associated protein 2 (MAP2) antibody (17490-1-AP, proteintech, China; 1:500). The corresponding secondary antibodies were used approximately, including Alexa Fluor 568-conjugated goat anti-human IgG antibody (A21090, ThermoFisher, USA; 1:1,000), and Alexa Fluor 488-conjugated goat anti-rabbit IgG antibody (A11008, ThermoFisher, USA; 1:1,000).

### Patient IgG preparation

CSF samples were obtained from patients with anti-NMDAR encephalitis, where anti-NMDAR antibodies had been verified by CBA. Total IgG was purified from CSF using the Melon Gel IgG Spin Purification Kit (45206, ThermoFisher, USA). Purified IgG was aliquoted and stored at –80 °C until use. This human-sample study was approved by the Ethics Committee of Nanfang Hospital, Southern Medical University (approval No. NFEC-2026-242).

### Passive-transfer mouse model (PTM)

Mice were anesthetized with isoflurane and secured in a stereotaxic frame. A 2-mm cranial skin incision was made, and lateral-ventricle injections were performed at the following coordinates: AP −0.22 mm, ML ±0.95 mm, DV −2.3 mm. Each mouse received 5 µl patient CSF-derived anti-NMDAR IgG (diluted to 80 µg/ml in PBS). dCLV ligation was carried out one-week prior to antibody administration. Animals were sacrificed 24 h after injection, and brain tissues were collected for analyses.

### Deep cervical lymph vessels (dCLV) ligation

All surgical operations were performed under anesthesia induced by intraperitoneal injection of ketamine (100 mg/kg) combined with xylazine (10 mg/kg). dCLV ligation was performed following previously reported procedures[24]. Briefly, the cervical skin was shaved and disinfected with iodine solution followed by 70 % ethanol. Ophthalmic ointment was applied to prevent corneal drying. A midline incision was made 5 mm above the clavicle. The sternocleidomastoid muscle was retracted to expose dCLV. Bilateral afferent lymphatic vessels leading to deep cervical lymph nodes (dCLN) were ligated at two sites, 0.1 mm and 0.5 mm proximal to each dCLN, using 11-0 non-absorbable nylon sutures. Sham-operated mice underwent identical surgical exposure without vessel ligation.

### Behavioral tests

All behavioral assessments were performed 12 weeks after the initial priming immunization by an investigator blinded to experimental group assignments. Behavioral outputs were recorded using the tracking system (TopScan 3.0, Clever Sys, USA).

#### Open-field test

Mice were acclimated to the testing room for at least one hour before testing. Each mouse was placed in the center of a 40 × 40 cm acrylic arena (cleaned with 75 % ethanol between subjects), facing away from the experimenter. Total travel distance, time spent in the center zone, and center-zone entry counts were recorded over a 5-min testing period.

#### Novel object recognition test

The test consisted of habituation, familiarization (T1), and test (T2) phases. During T1, two identical metal cubes (5 cm) were positioned symmetrically within the arena. Object exploration was defined as snout proximity ≤ 1 cm or direct paw contact lasting ≥ 1 s, and was recorded for 5 min. After a 1-hour interval, one metal cube was replaced with a novel cone-shaped object of matched volume (T2). Exploration durations toward the familiar (F) and novel (N) objects were used to calculate the discrimination index: DI = (N − F)/(N + F) × 100%. Arenas and objects were cleaned with 75% ethanol between individual trials.

#### Y-maze

The Y-maze test contained training and testing phases separated by a 1 h rest interval. During training, the novel arm was blocked by a removable guillotine-style door. Each mouse was placed in the start arm and allowed to freely explore the start and one open-arm for 10 min. Mice were then returned to their home cages. One hour later, the test phase was initiated: the novel arm was opened, and animals were placed in the start arm to explore all three arms for 5 min. Time spent in each arm and arm-entry events were quantified from recorded video during this 5 min session.

#### Elevated-plus maze test

The maze apparatus included two open-arms (36 cm × 6 cm) and two enclosed closed-arms (36 cm × 6 cm × 15 cm height), connected via a central 6 × 6 cm platform. The entire maze was elevated 50-60 cm above the floor. At 60 min after drug delivery, each mouse was placed inside a 60 × 60 × 35 cm plastic box for 5 min of environmental acclimation before being transferred onto the maze central platform. Open-arm and closed-arm entry counts and dwell time were recorded over 5 min. Percentages of open-arm entries (OE%) and open-arm time (OT%) were calculated relative to total arm entries and total time spent within arms.

#### Morris water maze (MWM)

A circular pool (120 cm diameter, water temperature 23-26°C) was divided into four virtual quadrants; water was made opaque using non-toxic white dye. A 9-cm hidden escape platform was submerged 1 cm below the water surface in the target (north) quadrant. Mice received four daily training trials with a 20-min inter-trial gap across five consecutive days. Release positions (east, west, north, south) were pseudo-randomized for each trial. If an animal failed to locate the platform within 60 s, it was gently guided onto the platform and permitted to remain there for 15 s. Average escape latency was calculated per mouse each day. On day 6, the platform was removed. Mice were released from the quadrant opposite to the target zone and tracked for 60 s. Time spent in the target quadrant and target-quadrant entry counts served as readouts for spatial-memory performance.

### Glymphatic influx evaluation

Anesthetized mice were positioned with the head flexed to ∼120°. After surgical exposure of the cisterna magna, a tracer mixture containing 8 µl tetramethylrhodamine-dextran (TMR-dextran, 70 kDa, 1 % w/v in artificial CSF; D1818, ThermoFisher, USA) plus 2 µl Alexa Fluor 488-conjugated goat anti-rabbit IgG antibody (rIgG-AF488) (2 mg/ml; A11008, ThermoFisher, USA) was injected at 3.125 µl/min using a PE-10 catheter connected to a 30-G needle. The needle remained in place for 60 s following infusion to prevent tracer reflux. Real-time transcranial imaging was performed on a stereomicroscope (M205 FA, Leica, Germany), capturing frames every 60 s over 60 min. Brains were subsequently harvested, fixed overnight in 4 % PFA at 4°C, and sectioned to 100 µm thickness with a vibratome (VT1200s, Leica, Germany). For each animal, 8 anatomically matched coronal slices were selected, and mean tracer fluorescence intensity was quantified with Image J software (NIH).

### Glymphatic efflux evaluation

Mice were anesthetized and a 0.5-mm burr hole was drilled over left (AP +1.5 mm, ML -1.5 mm, DV -2.5 mm) and right (AP +1.5 mm, ML +1.5 mm, DV -2.5 mm) parietal cortex. Alexa Fluor 555-labeled ovalbumin (OVA-555, 0.5 mg/ml in artificial CSF; O34782, ThermoFisher, USA) / and rIgG-AF488 were stereotactically injected into left and right thalamus respectively: 2 µl volume at an infusion rate of 0.2 µl/min. Four hours after circulation, brains were fixed, vibratome-sectioned (100 µm), and imaged on the stereomicroscope. Eight matched coronal slices per animal were analyzed to determine the fraction of area positive for OVA-555 and rIgG-AF488.

### AQP4 polarization analysis

To label cerebral vasculature, mice received tail-vein injection of 2 µl Dylight 488-conjugated tomato lectin (lectin-AF488; L32470, ThermoFisher, USA; 0.5 mg/ml). After 3-5 min circulation, animals were deeply anesthetized via intra-peritoneal sodium pentobarbital (50 mg/kg) and perfused transcardially with PBS followed by 4 % PFA. Coronal brain slices (100 µm) were permeabilized in 0.3 % Triton X-100 (15 min, room temperature) and blocked with 5 % BSA for 60 min. Sections were incubated overnight at 4°C with primary antibodies: rabbit anti-AQP4 antibody (ab125049, Abcam, USA; 1:200) and guinea-pig anti-GFAP antibody (173308, SYSY, Germany; 1:2,000). After PBS washes, sections were incubated with secondary antibodies Alexa Fluor 594-conjugated goat anti-rabbit IgG antibody (ab7149, Abcam, USA; 1:200) and Alexa Fluor 647-conjugated goat anti-guinea-pig IgG antibody (ab150187, Abcam, USA; 1:200) for 90 min at 37°C. Confocal images were acquired on a Zeiss LSM 980 microscope. The AQP4 polarization index was calculated as previously described: (peak perivascular AQP4 fluorescence − mean baseline fluorescence measured 10-20 µm away from vessels) / peak perivascular fluorescence [10, 25].

### Transmission electron microscope (TEM)

The mouse hippocampus tissue was dissected and fixed in electron-microscopy fixative (R20515, YUANYE, China) and processed through standard processing steps including dehydration, osmication, embedding, sectioning, and staining. Ultrastructural features of hippocampus were subsequently examined with a Hitachi H-7500 transmission electron microscope.

### Western blot

Mouse hippocampus tissue was lysed with denaturing lysis buffer (P0038, Beyotime, China) supplemented with protease inhibitors (FD1001, Fdbio, China) and phosphatase inhibitors (FD1002, Fdbio, China). Protein concentration was determined by BCA assay kit (FD2001, Fdbio, China). After boiled with loading buffer, 50 μg of protein was loaded per sample. Proteins were separated by 10% SDS–PAGE and subsequently transferred onto PVDF membranes. For NMDAR, membrane-enriched protein fractions were extracted from hippocampus using a commercial kit (BB3103, Bestbio, China) according to the manufacturer’s instructions. The isolated membrane proteins were mixed with loading buffer and denatured at 65 for 30 min. The membranes were probed with primary antibodies targeting GluN1 (27676-1-AP, Proteintech, China; 1:2,000), AQP4 (16473-1-AP, Proteintech, China; 1:2,000), β-dystroglycan (11017-1-AP, Proteintech, China; 1:2,000), Occludin (27260-1-AP, Proteintech, China; 1:2,000), α-Tubulin (11224-1-AP, Proteintech, China; 1:2,000) and GAPDH (10494-1-AP, Proteintech, China; 1:2,000), respectively. Band intensity was quantified using Image J software (NIH).

### IgG deposition

To determine the antibody deposition in the brain, floating brain slices were incubated with Alexa Fluor 488-conjugated goat anti-mouse IgG antibody (AB150113, abcam, USA; 1:200) or goat anti-human IgG antibody (A11013, ThermoFisher, USA; 1:200) for two days at 4 in PBS containing 0.03 % Triton-X100 and 3 % BSA. After washing, the images were taken by using a laser confocal microscope (LSM 980, Zeiss, Germany).

### Statistical analysis

Statistical analyses were performed using GraphPad Prism (version 10.3.1). The Shapiro-Wilk test was used to assess normality of data distribution. Parametric data were analyzed by Student’s t-test or one-way ANOVA followed by Tukey’s post-hoc test. Non-parametric data were analyzed with Mann–Whitney U test or Kruskal–Wallis tests. Statistical significance threshold was set at *P* < 0.05. Data were presented as mean ± SD. Graphs were prepared with GraphPad Prism and Adobe Illustrator CC 2024.

## RESULTS

### Establishment and validation of an active-immunization mouse model of anti-NMDAR encephalitis

The active-immunization mouse model of anti-NMDAR encephalitis was generated with polypeptides GluN1_356-385_ according to the protocol shown in **Fig. 1A**. Serum samples were collected 2, 4, 6, 8, 10, and 12 weeks after the initial immunization, and CSF was collected at week 12. Antibody titers were measured by a CBA assay using 293T cells overexpressing GluN1-eGFP fusion protein. GluN1-specific antibodies became detectable by week 2 with titers above 1:2,000. Titers rose after booster immunizations, reached a plateau at approximately week 10, and remained at 1:16,000 or higher (**Fig. 1B, C**). At week 12, the antibodies were detected in CSF (**Fig. 1 D**). No anti-NMDAR antibodies were detected in Sham mice. In TBA assay, serum from anti-NMDAR encephalitis mice labeled hippocampus neuropil and cerebellar granule cell layers in brain sections from naïve mice. This pattern resembled that produced by CSF from patients with anti-NMDAR encephalitis, whereas serum from Sham mice yielded no specific signal (**Fig. 1E**). When applied to primary rat hippocampus neurons cultured for 10-14 days in vitro, serum from anti-NMDAR encephalitis mice produced dendritic spine staining comparable to patient-derived IgG (**Supplementary Fig. S1**). Iba1 immunofluorescence was higher in the cortex and hippocampus CA1 and CA3 regions of anti-NMDAR encephalitis mice than in Sham mice (**Supplementary Fig. S2A, B**). Microglia in the cortex and hippocampus also showed reduced morphological complexity. Both branching index and maximal branch length decreased in these brain regions. Branch number remained unchanged in the cortex but reduced in the hippocampus (**Supplementary Fig. S2C, D**). These morphological changes are consistent with microglial activation. Behavioral tests revealed reduced total travel distance and shorter center-zone duration in the open-field test, a lower novel-object recognition index, and prolonged escape latency in the Morris water maze (**Fig. 1F**). These phenotypes indicate reduced locomotor activity, increased anxiety-like behaviors, and impaired recognition and spatial memory. Together, immunization with GluN1_356-385_ induced GluN1-reactive antibodies in serum and CSF, triggered microglial activation, and reproduced behavioral abnormalities characteristic of anti-NMDAR encephalitis.

### Glymphatic influx is enhanced in anti-NMDAR encephalitis mice

To assess influx function of GS in anti-NMDAR encephalitis, fluorescent tracers were injected into the cisterna magna (**Fig. 2A**). Influx was monitored by real-time transcranial macroscopic imaging. Brains were then harvested, sectioned, and imaged with fluorescence stereomicroscopy to quantify parenchymal tracer penetration. Aftercisterna magna TMR-dextran injection, tracer signal gradually accumulated along middle cerebral arteries and within olfactory bulbs. Tracer appeared earlier in anti-NMDAR encephalitis mice compared with Blank and Sham groups (10 min versus 20 min) and reached higher signal intensity at the 60 min time point (**Fig. 2B, C**). ex-vivo brain sections confirmed greater accumulation of TMR-dextran in anti-NMDAR encephalitis mice (**Fig. 2D, E**). We next assessed the influx efficiency for antibodies by injection of rIgG-AF488. Tracer signal appeared earlier in anti-NMDAR encephalitis mice, and the 60 min tracer positive area was larger than in Sham animals (**Fig. 2F, G**). Direct immunostaining of sagittal brain sections with fluorescence anti-mouse IgG secondary antibody showed higher endogenous IgG signal in anti-NMDAR encephalitis mice (**Fig. 2H, I**). Immunoblot analysis showed reduced NMDAR abundance in hippocampus of anti-NMDAR encephalitis mice compared with Sham controls (**Fig. 2J, K**). These data demonstrate enhanced influx of CSF-delivered solutes in anti-NMDAR encephalitis mice. This phenotype was accompanied by increased brain IgG deposition and reduced hippocampal NMDAR level.

**Figure 2.**
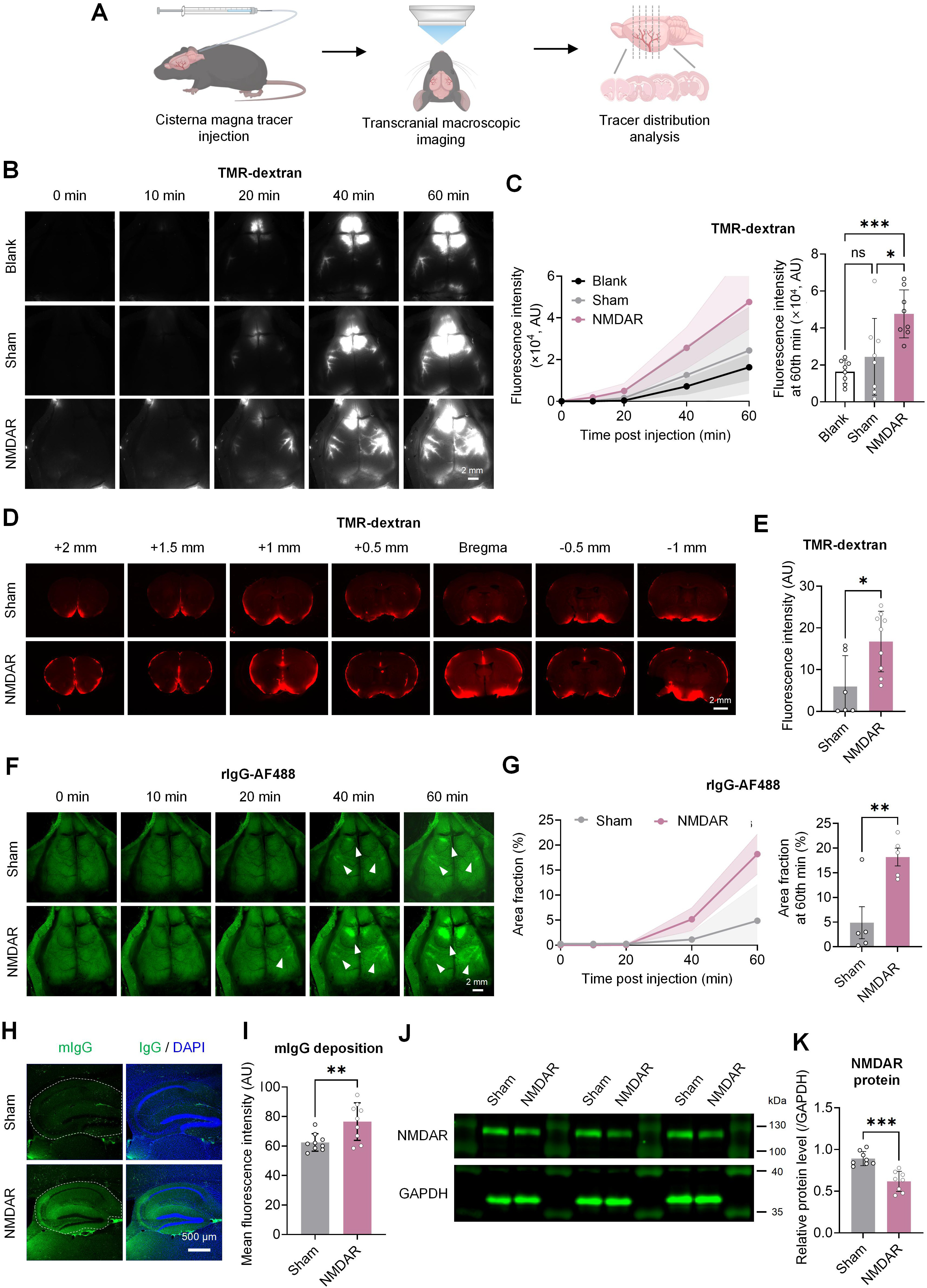
Glymphatic influx is enhanced in anti-NMDAR encephalitis mice, accompanied by increased brain IgG deposition and reduced NMDAR level. (A) Schematic workflow for evaluating influx function of glymphatic system. (B) Representative serial in vivo transcranial fluorescence images acquired at 0, 10, 20, 40, and 60 min after cisterna magna injection of TMR-dextran. Scale bar, 2 mm. (C) Time-course (left) and 60-min endpoint quantification (right) of TMR-dextran fluorescence intensity (Blank, n = 8; Sham, n = 7; NMDAR, n = 9). Untreated naïve mice served as blank controls. (D) Representative ex-vivo coronal brain sections showing the distribution of TMR-dextran across bregma from +2.0 to −1.0 mm. Scale bar, 2 mm. (E) Quantification of TMR-dextran fluorescence intensity in brain sections (Sham, n = 6; NMDAR, n = 9). (F) Representative serial fluorescence images obtained after cisterna magna injection of rIgG-AF488. Scale bar, 2 mm. (G) Time-course (left) and endpoint quantification (right) of rIgG-AF488-positive area (Sham, n = 7; NMDAR, n = 9). (H) Representative immunofluorescence images showing endogenous mouse IgG deposition in brain sections. Scale bar, 2 mm. (I) Quantification of the IgG fluorescence intensity (Sham, n = 9; NMDAR, n = 9). (J, K) Representative immunoblots (J) and quantitative analysis (K) of NMDAR protein level in hippocampus (Sham, n = 8; NMDAR, n = 8). rIgG-AF488, Alexa Fluor 488-labelled goat anti-rabbit IgG antibody; TMR-dextran, tetramethylrhodamine-labelled dextran, 70 kDa. \**P* < 0.05, \*\**P* < 0.01, \*\*\**P* < 0.001, \*\*\*\**P* < 0.0001; ns, not significant.

### Perivascular AQP4 polarization is increased in anti-NMDAR encephalitis mice

Perivascular localization of AQP4 on astrocytic endfeet is essential for efficient glymphatic transport. We therefore analyzed AQP4 polarization and astrocyte reactivity in hippocampus sections. Blood vessels and astrocytes were labeled with lectin-AF488 and GFAP, respectively. Global AQP4 fluorescence intensity showed no obvious difference between anti-NMDAR encephalitis and Sham mice. In contrast, the AQP4 polarization index was higher in anti-NMDAR encephalitis mice (**Fig. 3A-C**). Elevated GFAP fluorescence intensity and larger GFAP positive area indicated astrocyte activation in anti-NMDAR encephalitis mice (**Fig. 3D, E**). Western blot results showed comparable total protein levels of AQP4, β-DG and occludin across groups (**Fig. 3F, G**). TEM analysis showed intact BBB tight-junctions, whereas astrocytic endfeet exhibited swelling (**Fig. 3H**). These observations point toward AQP4 redistribution toward perivascular astrocytic endfeet, rather than elevated total AQP4 protein expression.

**Figure 3.**
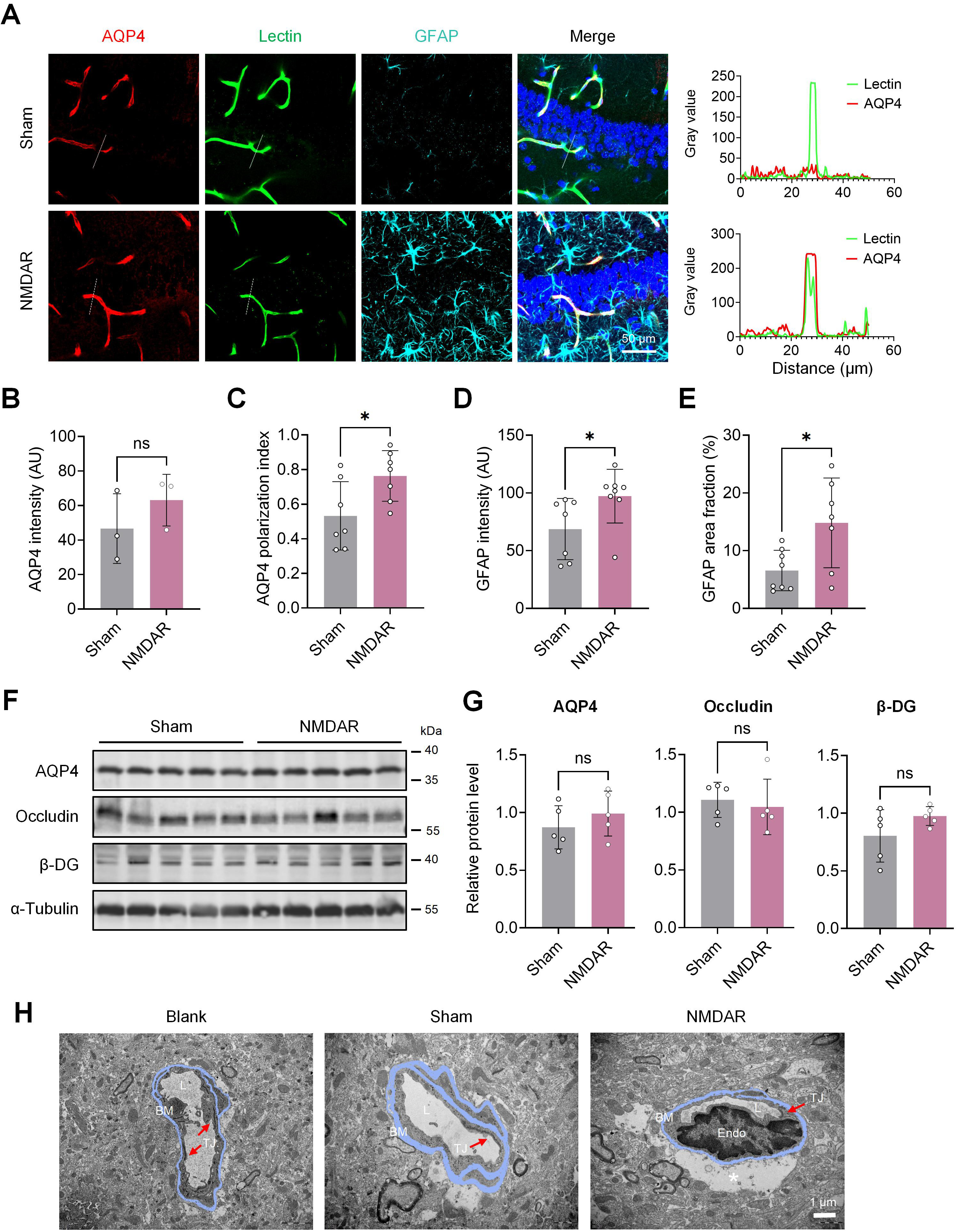
Perivascular AQP4 polarization is increased in anti-NMDAR encephalitis mice. (A) Representative immunofluorescence images of brain sections. AQP4 is shown in red, lectin-labeled blood vessels in green, GFAP in cyan, and nuclear DAPI staining in blue. Scale bar, 50 μm. Distribution of AQP4 and Lectin labeled blood vessels is shown in right panels. Line panels on the right illustrate spatial distribution of AQP4 relative to lectin-positive vessels. (B) Quantification of AQP4 fluorescence intensity (Sham, n = 3; NMDAR, n = 3). (C) Quantification of the AQP4 polarization index. The index was calculated as: (peak perivascular AQP4 fluorescence - mean baseline fluorescence measured 10-20 µm away from vessels) divided by peak perivascular fluorescence (Sham, n = 7; NMDAR, n = 7). (D) Quantification of GFAP fluorescence intensity (Sham, n = 6; NMDAR, n = 8). (E) Quantification of GFAP-positive area fraction (Sham, n = 8; NMDAR, n = 7). (F) Western blotting of AQP4, Occludin, and β-DG proteins in the hippocampus. (G) Quantification of relative protein levels of AQP4, Occludin, and β-DG. (H) Representative transmission electron microscopy images of the blood-brain barrier in hippocampus. BM, basement membrane; Endo, vascular endothelial cell; L, lumen; TJ, tight junction (red arrows); asterisk in right picture indicates astrocytic endfoot. Scale bar, 1 μm. \**P* < 0.05; ns, not significant.

### Inhibition of AQP4 suppresses glymphatic influx and improves behavior phenotypes in anti-NMDAR encephalitis mice

To determine whether GS mediates antibody entry into the brain parenchyma, we administered AER-271, a selective AQP4 inhibitor, to interfere with glymphatic function. Mice received intraperitoneal AER-271 (5 mg/kg) every two days for 7 consecutive days starting at week 11 (**Fig. 1A**). After treatment, behavioral tests and tracer imaging experiments were performed.

In the open-field test, anti-NMDAR encephalitis mice treated with AER-271 showed recovery in total travel distance and time spent in the central zone, whereas vehicle treated NMDAR animals did not display such improvement (**Fig. 4A, B**). These changes indicate improved locomotor and center-directed exploratory behavior. TMR-dextran and rIgG-AF488 were co-injected into the cisterna magna. AER-271 delayed the initial appearance of both tracers and reduced their fluorescence signals during real-time transcranial imaging (**Fig. 4C, D**). Ex-vivo brain section analysis also showed lower TMR-dextran accumulation after AER-271 treatment (**Supplementary Fig. S3**).

**Figure 4.**
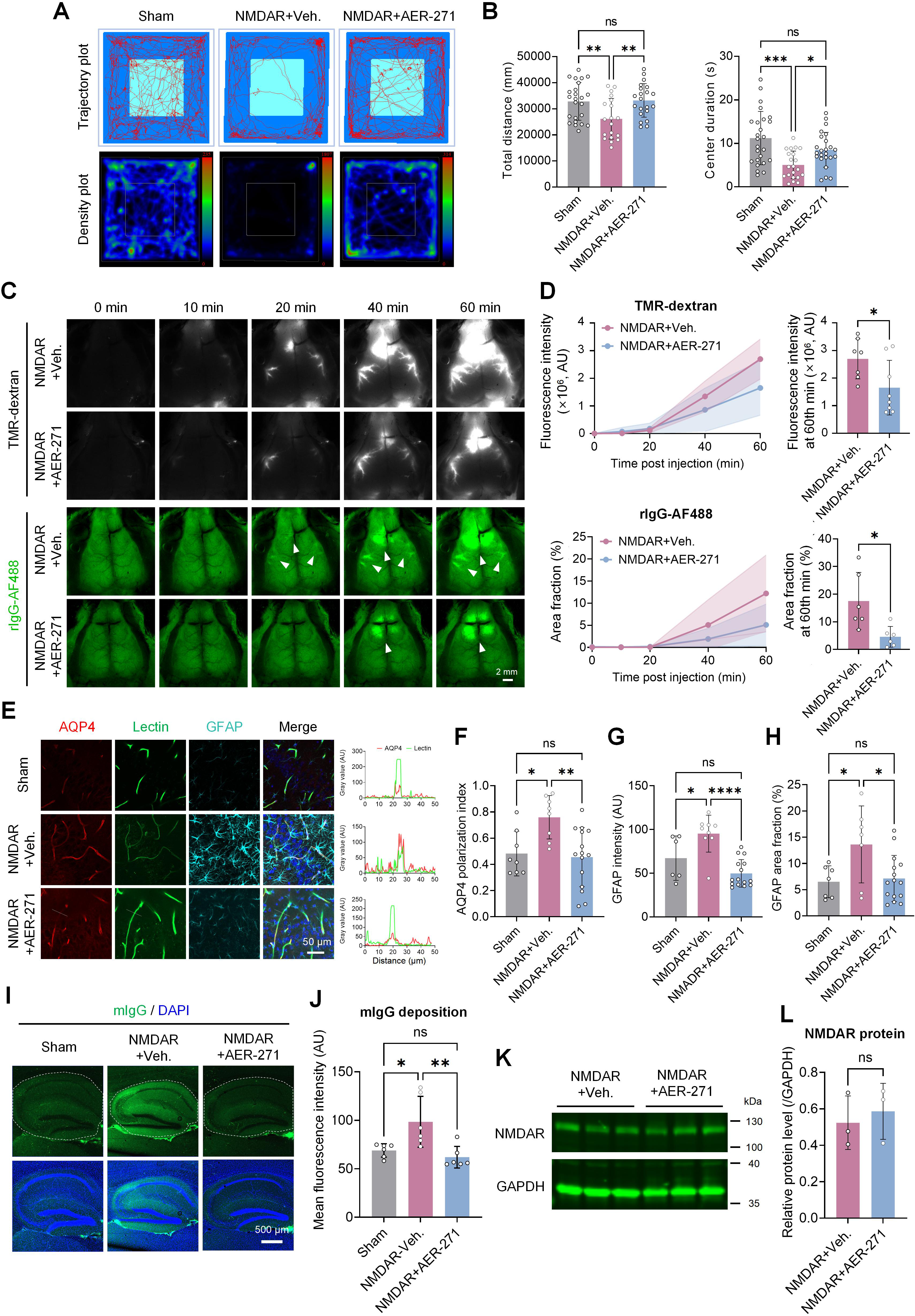
AQP4 inhibition with AER-271 reduced glymphatic influx, AQP4 polarization, and IgG deposition, and restored behavioral phenotypes in anti-NMDAR encephalitis mice. (A) Representative open-field test trajectory plots and occupancy density heat maps for Sham, NMDAR + vehicle (Veh.), and NMDAR + AER-271 groups. (B) Quantification of the total travel distance (Sham, n = 23; NMDAR + Veh., n = 18; NMDAR + AER-271, n = 22) and time spent within the central zone (Sham, n = 25; NMDAR + Veh., n = 21; NMDAR + AER-271, n = 24). (C) Representative serial fluorescence images acquired after the simultaneous cisterna magna injection of TMR-dextran (upper panels) and rIgG-AF488 (lower panels). Arrowheads indicate rIgG-AF488 tracer signal. Scale bar, 2 mm. (D) Time-course and 60-min endpoint quantification of TMR-dextran fluorescence intensity (NMDAR + Veh., n = 7; NMDAR + AER-271, n = 8) and the rIgG-AF488-positive area fraction (n = 6 per group). (E) Representative immunofluorescence images of brain sections. AQP4 is shown in red, lectin-AF488 in green, GFAP in cyan, and DAPI-stained nuclei in blue. Scale bar, 50 μm. (F) Quantification of AQP4 polarization index. (Sham, n = 8; NMDAR + Veh., n = 8; NMDAR + AER-271, n = 15). (G) Quantification of GFAP fluorescence intensity (Sham, n = 6; NMDAR + Veh., n = 9; NMDAR + AER-271, n = 15). (H) Quantification of GFAP-positive area fraction (Sham, n = 7; NMDAR + Veh., n = 7; NMDAR + AER-271, n = 15). (I) Representative immunofluorescence images showing endogenous mouse IgG (mIgG, green) deposition within the hippocampus. Nuclei were counter-stained with DAPI (blue); dashed lines outline regions selected for quantitative analysis. Scale bar, 500 μm. (J) Quantification of fluorescence intensity of mIgG in hippocampus (n = 6 per group). (K, L) Representative immunoblots (K) and quantification (L) of NMDAR protein in hippocampus (n = 3 per group). \**P* < 0.05, \*\**P* < 0.01, \*\*\**P* < 0.001, **** *P* < 0.0001; ns, not significant.

Consistent with earlier observations, the hippocampus exhibited higher AQP4 polarization in anti-NMDAR encephalitis mice relative to Sham controls, and this elevated polarization was significantly reduced after AER 271 treatment (**Fig. 3E–H**). Hippocampus IgG deposition was reduced in AER-271-treated mice compared with vehicle-treated anti-NMDAR encephalitis mice (**Fig. 4I, J**). However, hippocampus NMDAR abundance showed no significant difference between two groups (**Fig. 4K, L**). Thus, AER-271 reduced glymphatic influx, lowered excessive AQP4 polarization, and reduced hippocampal IgG deposition, which in turn partially alleviated disease-related behavioral deficits in anti-NMDAR encephalitis mice. These effects occurred without detectable recovery of NMDAR protein level.

### Glymphatic efflux is enhanced in anti-NMDAR encephalitis mice

We next set out to determine whether efflux function of GS was also changed by measuring the clearance of tracers directly injected into the brain parenchyma. Two hours after intra-parenchymal tracer injection, anti-NMDAR encephalitis mice retained less OVA-555 signal in brain tissue compared with Sham animals (**Supplementary Fig. S4**). Further, we stereotactically injected OVA-555 and rIgG-AF488 into the left and right thalamus, respectively (**Fig. 5A**). Four hours later, anti-NMDAR encephalitis mice retained less OVA-555 and rIgG-AF488 within brain than Sham mice (**Fig. 5B, C**). Compared with vehicle treatment, AER 271 administration increased residual tracer retention for both probes. Consistent with these brain clearance results, OVA-555 and rIgG-AF488 fluorescence signals in dCLN decreased after AER-271 treatment (**Fig. 5D**). These data indicate that glymphatic efflux is enhanced in anti-NMDAR encephalitis mice, and this enhanced efflux can be blunted by pharmacological AQP4 inhibition.

**Figure 5.**
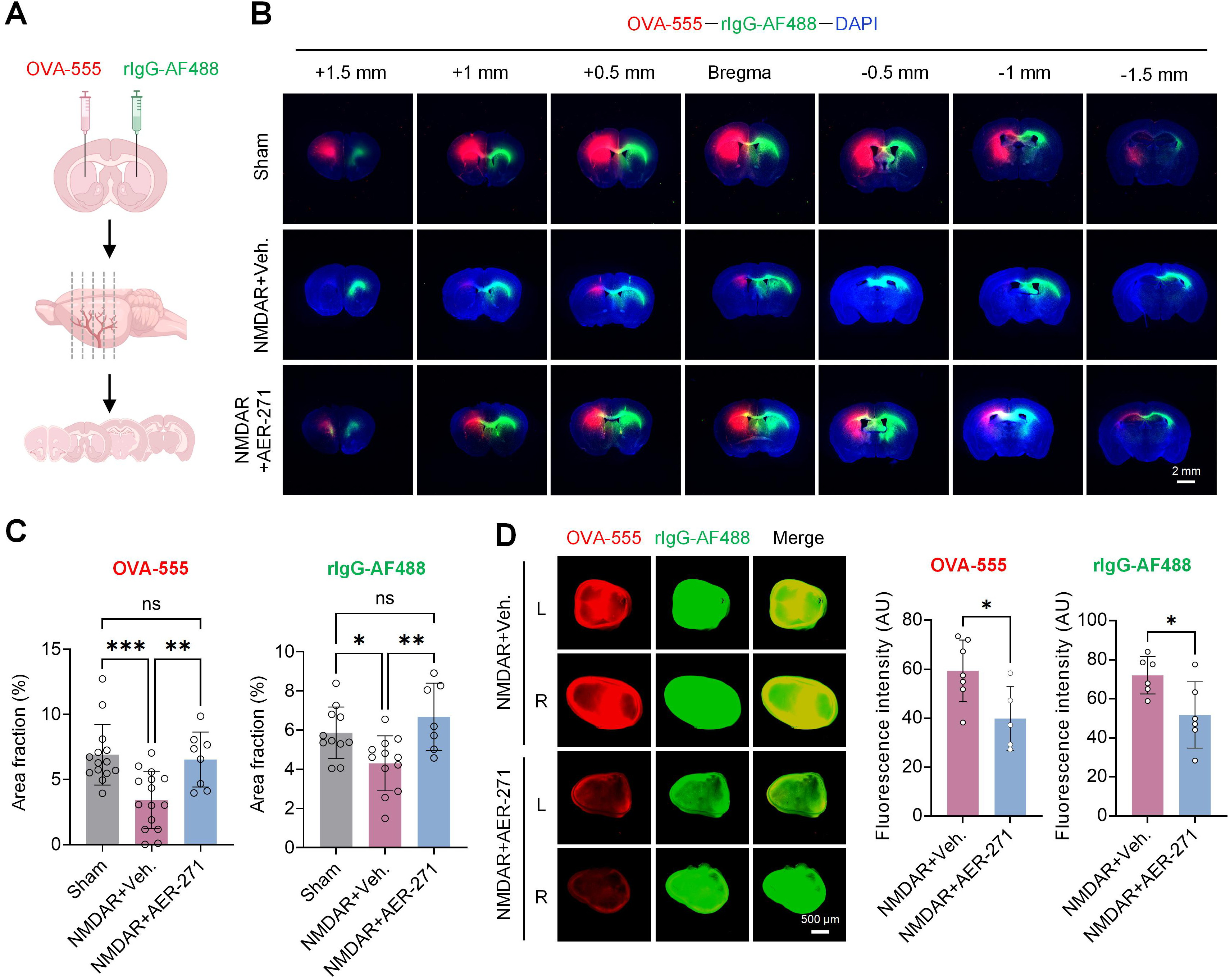
AER-271 attenuates enhanced glymphatic clearance in anti-NMDAR encephalitis mice. (A) Schematic workflow for assessing efflux or clearance function of glymphatic system. OVA-555 and rIgG-AF488 were stereotactically injected into the left and right thalamus, respectively. Brains were harvested 4 h post-injection, sectioned, and residual tracer signals were quantified to evaluate clearance efficiency. (B) Representative coronal brain sections (bregma +1.5 mm to -1.5 mm) imaged 4 h after tracer injection. OVA-555 signal in red; rIgG-AF488 in green; DAPI nuclear staining in blue. Scale bar, 2 mm. (C) Quantification of the tracer-positive area fraction across the analyzed brain sections For OVA-555: Sham, n = 15; NMDAR + Veh., n = 15; NMDAR + AER-271, n = 8. For rIgG-AF488: Sham, n = 11; NMDAR + Veh., n = 12; NMDAR + AER-271, n = 7. (D) Representative fluorescence images and quantitative analysis for OVA-555 and rIgG-AF488 drainage into dCLNs. L and R denote left and right dCLNs. OVA-555: NMDAR + Veh., n = 7; NMDAR + AER-271, n = 5. rIgG-AF488: NMDAR + Veh., n = 6; NMDAR + AER-271, n = 6. Scale bar, 500 µm. dCLNs, deep cervical lymph nodes; OVA-555, Alexa Fluor 555-labeled ovalbumin; rIgG-AF488, Alexa Fluor 488-labelled goat anti-rabbit IgG antibody. \**P* < 0.05, \*\**P* < 0.01, and \*\*\**P* < 0.001; ns, not significant.

### Ligation of dCLV impairs antibody clearance and increases anxiety-like behavior in a passive-transfer model

To test the role of extracranial lymphatic drainage, we performed bilateral upstream dCLV ligation in the passive-transfer model (PTM) of anti-NMDAR encephalitis. Surgical ligation was performed 7 days before intracerebroventricular injection of patient-derived anti-NMDAR IgG. Behavior tests and tissue analyses were conducted 24 h after antibody administration (**Fig. 6A**). Surgical efficacy was validated by cisterna magna Evans blue infusion. Dye readily accumulated within dCLNs in sham-operated animals, yet was absent from these lymph nodes after successful dCLV ligation (**Fig. 6B**).

**Figure 6.**
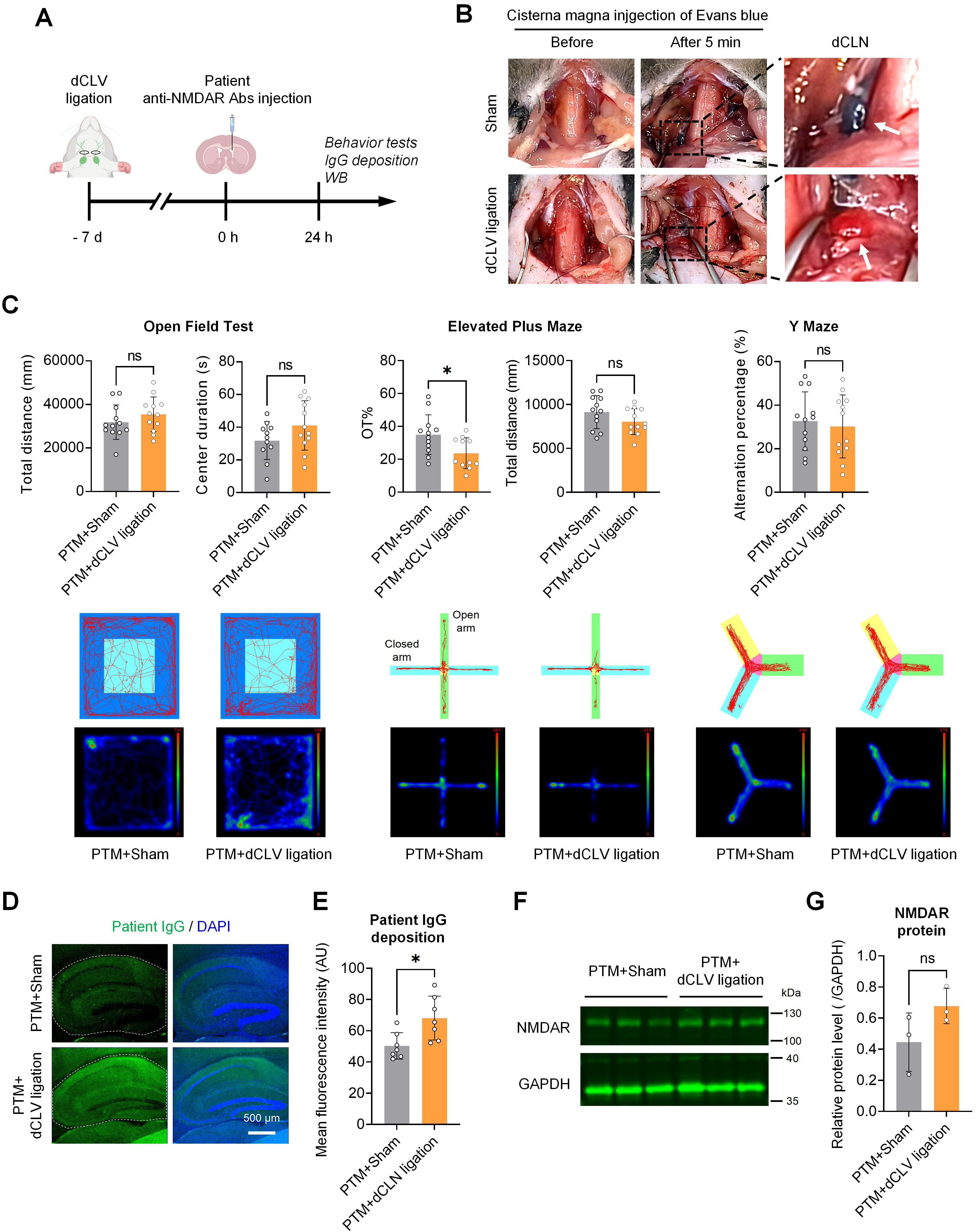
Ligation of deep cervical lymphatic vessel increases brain retention of patient-derived anti-NMDAR IgG and exacerbates anxiety-like behavior in the passive-transfer model. (A) Experimental workflow. dCLV ligation (proximal to dCLN) was performed seven days before passive antibody transfer. Patient CSF derived anti NMDAR IgG was injected into the lateral ventricle to establish the passive transfer model (PTM + dCLV ligation). Behavioral testing and tissue analysis were performed 24 h after intraventricular antibody delivery. PTM mice receiving identical surgical exposure without ligation served as PTM Sham controls. (B) Validation of dCLV ligation. Following intraventricular Evans-blue injection, dye accumulated within dCLNs in sham-operated animals, but was absent after successful dCLV ligation. White arrows indicate the dCLN. (C) Representative movement trajectories and occupancy heat maps (lower panels), together with quantitative behavioral analyses (upper panels). Open-field test: total distance, PTM + Sham n = 12, PTM + dCLV ligation n = 12; central zone duration, n = 11 and 12 respectively. Elevated-plus maze: open-arm dwell time, n = 12 and 11 respectively; total travel distance n = 12 for each group. Y-maze spontaneous alternation test, n = 12 per group. (D, E) Representative fluorescence images (D) and quantitative analysis (E) of patient-derived IgG deposition within the hippocampus. Patient-derived IgG signal is shown in green; DAPI nuclear staining in blue. Dashed lines outline the analyzed hippocampus region. PTM + Sham, n = 7; PTM + dCLV ligation, n = 7. Scale bar, 500 μm. (F, G) Representative immunoblots (F) and quantification (G) of NMDAR protein in hippocampus (n = 3 mice per group). NMDAR levels normalized to GAPDH showed no significant difference between groups. dCLN, deep cervical lymph nodes; dCLV, deep cervical lymphatic vessel; PTM, passive-transfer model. \**P* < 0.05; ns, not significant.

In the open-field and Y-maze tests, we observed no significant behavioral differences between ligated and sham-operated PTM groups. In the elevated-plus maze, however, dCLV-ligated PTM mice spent a smaller proportion of time in open-arms, while their overall locomotor activity remained unchanged (**Fig. 6C**). This finding indicates that dCLV ligation aggravated anxiety-like phenotypes without affecting general locomotion or working-memory performance. Patient-derived anti-NMDAR IgG in hippocampus was higher in ligated mice, consistent with impaired antibody clearance and increased brain retention (**Fig. 6D, E**). Hippocampus NMDAR level did not differ significantly between the two groups (**Fig. 6F, G**). Together, these data show that impaired downstream drainage of GS reduced the brain clearance of patient-derived IgG and exacerbated anxiety-like behavior in PTM mice. These results suggest that disruption of downstream glymphatic-lymphatic drainage diminishes brain antibody clearance and exacerbates anxiety-related behaviors in the PTM mice.

### Dexmedetomidine enhances brain antibody clearance and reduces IgG deposition

To examine whether pharmacological enhancement of glymphatic function could affect antibody clearance, anti-NMDAR encephalitis mice received subcutaneous Dex, which was dissolved in saline and dosed at 0.2 mg/kg every two days starting at week 11 for a total of seven days (**Fig. 1A**). In the open-field test, Dex-treated mice exhibited reduced total travel distance and shorter central-zone duration time relative to vehicle treated mice. No group differences were detected in elevated-plus maze and Y-maze tests (**Fig. 7A**). Two hours after intrathalamic tracer injection, brains from Dex treated mice displayed lower residual OVA-555 and rIgG-AF488 signals (**Fig. 7B, C**). In line with this improved tracer clearance, endogenous mouse IgG within the hippocampus were also decreased in the Dex-treated group (**Fig. 7D, E**). Therefore, dexmedetomidine boosted parenchymal clearance of antibody, accompanied by reduced hippocampal IgG deposition.

**Figure 7.**
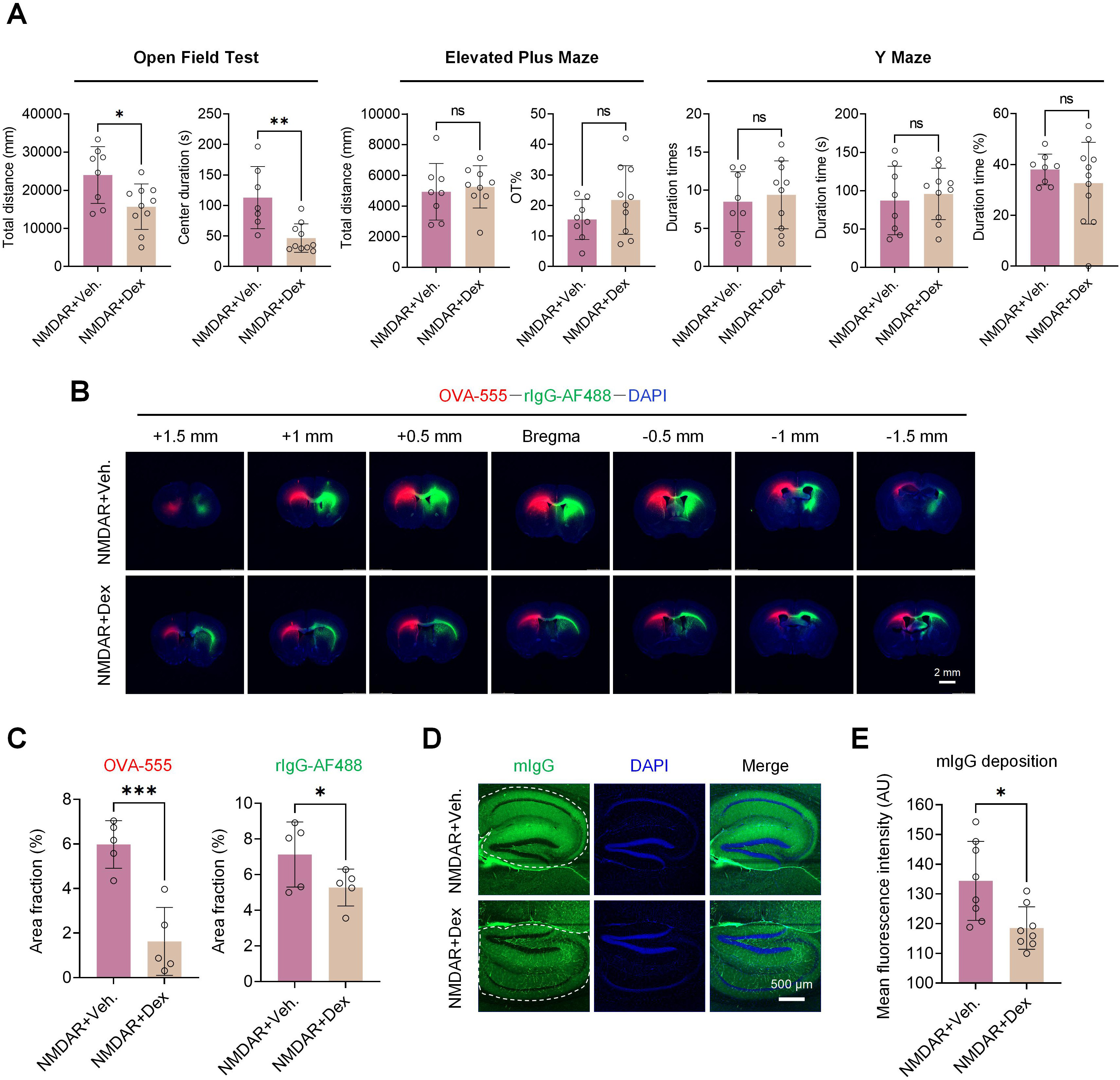
Dexmedetomidine enhances glymphatic efflux in anti-NMDAR encephalitis mice. (A) Behavioral assessments of anti-NMDAR encephalitis mice treated with dexmedetomidine (NMDAR + Dex) versus vehicle control (NMDAR + Veh.). Open-field test: total distance, NMDAR + Veh. n = 8, NMDAR + Dex n = 10; central zone duration, n = 7 and 10 respectively. Elevated-plus maze, n = 8 and 10 respectively. Y-maze: novel arm entry count, novel arm dwell time, and percentage time spent in the novel arm: NMDAR + Veh., n = 8, 8, 8; NMDAR + Dex, n = 10, 10, 11. (B) Representative coronal brain sections (bregma +1.5 mm to -1.5 mm), 4 h after stereotactic thalamic injection of OVA-555 and rIgG-AF488. OVA-555 is shown in red; rIgG-AF488 in green; DAPI in blue. Scale bar, 2 mm. (C) Quantification of the tracer-positive area across brain sections (n = 5). (D, E) Representative fluorescence images (D) and quantitative analysis (E) of endogenous mouse IgG deposition in the hippocampus (n = 8). Mouse IgG signal in green; DAPI nuclear staining in blue. Scale bar, 500 μm. \**P* < 0.05, \*\**P* < 0.01, \*\*\**P* < 0.001; ns, not significant.

## DISCUSSION

The present study provides novel evidence that the GS is dynamically upregulated in an active-immunization mouse model of anti-NMDAR encephalitis, and that this upregulation plays a complex, dual role in the disease. Our findings show that both glymphatic influx and efflux are enhanced in the diseased state, coupled with increased perivascular polarization of AQP4 on astrocytic endfeet. Pharmacological inhibition of the GS with AER-271, which reduces AQP4 polarization, attenuated both glymphatic influx and efflux, decreased brain deposition of endogenous IgG, and partially ameliorated behavioral deficits. Conversely, surgical ligation of the dCLVs, a major extracranial efflux route, impaired antibody clearance and exacerbated anxiety-like behavior. The schematic model (**Fig. 8**) integrates the anatomical architecture of the GS with the molecular pathogenesis of anti-NMDAR encephalitis. Under normal conditions, CSF enters the brain along periarterial spaces, driven by polarized AQP4 on astrocytic endfeet. In the diseased state, anti-NMDAR antibodies bind to neuronal NMDARs, leading to enhanced AQP4 phosphorylation and membrane polarization. The resulting hyperactive GS exerts a “double-edged sword” effect: accelerated influx facilitates further antibody entry from the subarachnoid space into the parenchyma, while enhanced efflux promotes antibody clearance via the perivenous space and meningeal lymph vessels toward deep cervical lymph nodes. These findings suggest that the GS is not merely a passive clearance pathway in anti-NMDAR encephalitis but an active participant in both the distribution and removal of pathogenic antibodies.

**Figure 8.**
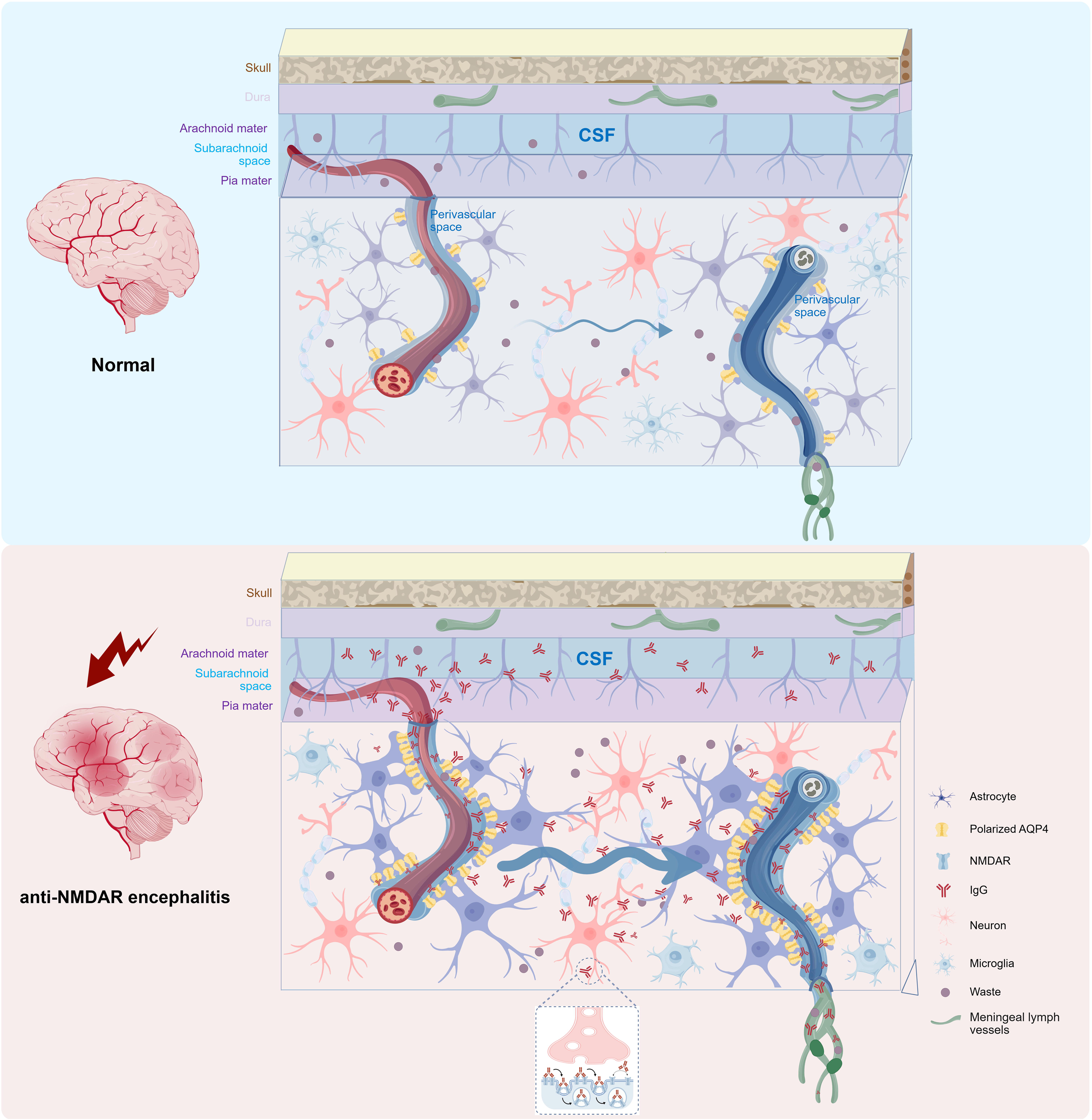
Proposed model for glymphatic system-driven antibody transport in the anti-NMDAR encephalitis brain. Under physiological conditions, perivascular astrocyte endfoot AQP4 supports CSF influx along periarterial spaces, facilitates CSF-ISF exchange within brain parenchyma, and enables waste clearance through perivenous drainage and meningeal lymphatic pathways. In anti-NMDAR encephalitis, elevated perivascular AQP4 polarization augments both glymphatic influx and efflux. Enhanced influx drives entry of CSF-borne anti-NMDAR IgG into brain parenchyma. These antibodies bind surface neuronal NMDAR and trigger receptor internalization, lowering total NMDAR abundance and producing neuronal dysfunction. Meanwhile, augmented glymphatic efflux promotes removal of IgG from brain tissue. (The graph is generated by using www.biogdp.com)

Contrary to the glymphatic suppression often reported in neurodegenerative conditions like Alzheimer’s disease [26, 27], we observed a pronounced enhancement of both glymphatic influx and efflux functions in our model. The accelerated parenchymal entry of intracisternally injected TMR-dextran and rIgG-AF488 antibody and the increased clearance of intrastriatally injected tracers (OVA-555 and rIgG-AF488) strongly suggest an overall facilitation of fluid transport through the brain’s interstitial spaces. This finding positions anti-NMDAR encephalitis as a unique neuroinflammatory condition characterized by hyperactive, rather than impaired, glymphatic function. This divergence suggests that the GS response is disease specific. In the context of active neuroinflammation, as seen in our model, the GS may be adaptively modulated. This is consistent with other studies showing that in certain pathological states, such as high-fat diet-induced obesity or decompression sickness, despite the presence of neuroinflammation, GS function can be enhanced, potentially as a compensatory mechanism to accelerate the clearance of toxins and inflammatory mediators [28, 29]. Our findings align with this concept, proposing that the glymphatic upregulation in anti-NMDAR encephalitis could be a double-edged sword: it may facilitate the clearance of antibodies and inflammatory cytokines, but it could also paradoxically increase the influx of these same pathogenic molecules from the CSF into the brain parenchyma, thereby exacerbating neuronal injury. We hypothesized that the net effect depends on the balance between inflow and outflow.

An intriguing finding of the present study is that the enhanced glymphatic function and antibody brain entry in anti-NMDAR encephalitis occurred in the absence of detectable BBB disruption. Our TEM analysis revealed intact tight junctions, and Western blotting showed no significant changes in the protein levels of AQP4, β-DG, or Occludin. These ultrastructural and molecular data are consistent with the clinical observation[4, 5] and that serum albumin-based measures of BBB integrity (QAlb) are normal in a substantial proportion of patients [5, 30]. Together, these observations challenge the traditional view that passive leakage across a disrupted BBB is the primary route for antibody entry into the brain parenchyma in this disease [3, 31].

The GS function is intimately dependent on AQP4, which is highly polarized to the perivascular endfeet of astrocytes and facilitates convective fluid transport[6, 32]. Our data demonstrate significant astrocyte activation, alongside an enhanced AQP4 polarization to perivascular endfeet. The increase in the polarization would theoretically lower the resistance for water flux between the perivascular space and the neuropil [33]. This reorganization of AQP4 is likely a primary driver of the observed glymphatic upregulation. The electron microscopy findings of astrocytic end-foot edema in the model group further support heightened fluid flux.

Our findings represent a distinct pathological state that differs from the classical stroke and TBI [34, 35]. In acute brain injuries such as ischemic stroke, AQP4 exhibits a well-documented biphasic, time-dependent pattern [36]. During the acute phase (hours to 24 h post-injury), AQP4 expression is rapidly upregulated and its membrane distribution at the perivascular endfeet is maintained or increased, which facilitates water influx into astrocytes, driving cytotoxic edema. At this early stage, AQP4 is detrimental. However, in the subacute to chronic phase (48 h to weeks), AQP4 undergoes progressive depolarization, leading to impaired glymphatic clearance and the transition to vasogenic edema [37]. At this later stage, the loss of AQP4 polarization exacerbates pathology by preventing the clearance of excess fluid and metabolic waste, and partial restoration of polarization during the resolution phase supports edema clearance[34, 38].

We propose that this reflects an intermediate phase of GS hyperactivation that has not yet transitioned to the destructive, depolarization-driven dysfunction seen in late-stage ischemia. The endfoot swelling observed by TEM may represent the earliest structural correlate of this hyperdynamic state, analogous to the acute phase of ischemic injury, but the preserved AQP4 polarization and patent efflux pathways (including meningeal lymphatics) may prevent the accumulation of excess fluid that would otherwise manifest as MRI-visible edema. This interpretation is consistent with the clinical observation that the majority of anti-NMDAR encephalitis patients present with normal MRI findings [4],

The trigger for this astrocytic response and AQP4 repolarization is unclear. It is likely the presence of anti-NMDAR antibodies themselves. NMDARs are expressed on astrocytes[39, 40], and their antagonism or modulation by anti-NMDAR autoantibodies could directly alter astrocytic calcium signaling, gene expression, and ultimately, their polarized phenotype[41]. Whether this represents a novel underlying mechanism whereby a specific autoantibody directly modulates the fundamental physiology of cerebral fluid transport remains to be determined.

The most intriguing finding is that inhibition of AQP4 with AER-271, while effectively suppressing the enhanced glymphatic function and normalizing AQP4 polarization, led to an improvement in behavioral deficits. The hyperactive GS might contribute to pathophysiology by facilitating an excessive and uncontrolled flux of autoantibodies and inflammatory cytokines from the CSF into the brain parenchyma. While intended for clearance, this heightened influx might inadvertently worsen the distribution and synaptic concentration of pathogenic antibodies, exacerbating NMDAR hypofunction [42]. By inhibiting AQP4 and slowing glymphatic influx, AER-271 may limit the parenchymal penetration of antibodies, thereby reducing their synaptic impact. However, in this study, it did not restore hippocampus NMDAR protein levels. This might be attributed to insufficient treatment duration for NMDAR recovery. The state of astrocyte activation and AQP4 polarization, while facilitating fluid flow, may represent a maladaptive response that disrupts other critical astrocytic functions, such as potassium buffering or glutamate reuptake [43]. Normalizing this activated state with AER-271 might restore a more balanced astrocytic profile.

The partial nature of the behavioral improvement—significant in the open-field test (anxiety-like behavior) but not in novel object recognition (memory)—suggests that while modulating glymphatic flux and astrocyte activation is beneficial, it does not address the root cause: the presence of the autoantibodies. The primary therapeutic goal in anti-NMDAR encephalitis remains immunomodulation to eliminate antibodies[44]. However, our findings position AQP4 and the GS as a potential druggable pathway for symptomatic and disease-modifying adjunct therapy.

The dCLV ligation experiment directly confirms the importance of the meningeal lymphatic system as the final common pathway for brain waste clearance [45, 46]. The observation that impaired drainage leads to increased brain IgG retention and exacerbated anxiety-like behavior is a pivotal finding. It not only underscores the role of the GS in antibody clearance but also provides a direct link between GS function and behavioral phenotypes. This is consistent with the growing body of evidence that the meningeal lymphatics are a critical interface between the CNS and the peripheral immune system, and their dysfunction can contribute to neuroinflammation and cognitive decline [47].

The pharmacological modulation of the GS with Dex in our anti-NMDAR encephalitis model yielded results that are both intriguing and complex. Dex, a known GS enhancer[18], accelerated the clearance of OVA-555 and rIgG-AF488, and reduced hippocampus endogenous IgG deposition, supporting the concept that augmenting GS efflux can facilitate the removal of molecules from the brain. However, it did not improve behavioral outcomes. In fact, Dex reduced spontaneous locomotion.

This paradox can be explained by Dex’s dual effect: while enhancing efflux, it also accelerates glymphatic influx [48]. In a disease like anti-NMDAR encephalitis, where antibody entry is a primary driver of pathology, this could simultaneously increase the delivery of new antibodies into the brain parenchyma, counteracting the benefits of enhanced clearance. Furthermore, the drug’s sedative properties confounded the behavioral readouts. These findings serve as a cautionary note: simply “upregulating” the entire GS is not a straightforward therapeutic strategy. The net effect depends on the balance between augmenting clearance (the “drain”) and inadvertently promoting the entry of pathogenic molecules (the “inlet”), suggesting that future therapies should aim for selective enhancement of the efflux pathway, or combined with elimination of pathogenic sources.

Though Dex enhanced the clearance, clearance efficiency was lower for rIgG-AF488 compared with OVA-555. This discrepancy suggests that the ability of Dex to enhance GS-mediated clearance may be size-dependent. Another unexpected result was that the reduction in hippocampus endogenous IgG deposition without a corresponding improvement in behavior. This finding suggests that the deposition measured at a single time point (24 hours after the last Dex dose) may not reflect the cumulative pathological burden. The behavioral deficits observed in this model are likely the result of a complex interplay of factors, including NMDAR internalization, synaptic dysfunction, neuroinflammation, and the structural damage caused by astrocyte endfoot swelling [49]. Simply clearing some IgG at a late stage may be insufficient to reverse these established pathological processes, especially if the enhanced influx simultaneously introduces new antibodies and inflammatory mediators. The time course of the intervention may also be critical; a short-term, acute administration of Dex may not be sufficient to achieve a meaningful therapeutic effect, whereas a more sustained or early intervention might produce different results.

This research advances our understanding of the interplay between autoimmunity and brain fluid dynamics, highlighting the GS as an important player in neuroinflammatory disease and a promising therapeutic target. However, this study has several limitations. The active-immunization mouse model does not fully recapitulate the severe clinical features of human anti-NMDAR encephalitis, such as seizures, autonomic abnormalities. Antibody clearance was assessed at a single time point, ignoring the known diurnal variation of glymphatic function. The molecular mechanisms driving enhanced AQP4 polarization were not causally validated. Future studies should employ more disease models, longitudinal imaging, mechanism digging, diverse interventions, and clinical evidence to address these gaps.

### Conclusions

In summary, our study describes a previously unrecognized facet of anti-NMDAR encephalitis pathophysiology: a disease-associated glymphatic upregulation driven by enhanced astrocytic AQP4 polarization. This is unlike the conventional view that neuroinflammation always impairs glymphatic clearance, suggesting a more nuanced, disease-specific regulation. The beneficial effect of AQP4 inhibition opens a new avenue for therapeutic exploration. Instead of broadly boosting glymphatic activity, precise tuning of glymphatic transport may represent a viable strategy for treating autoimmune encephalitis.

## DECLARATIONS

### Availability of data and materials

The datasets of the current study are available from the corresponding author upon reasonable request.

### Authors’ contributions

Conceptualization: K.X., Y.W., D.W.; Data curation: J.L., F.H.; Formal analysis: J.L., F.H., K.X.; Funding acquisition: K.X., D.W., Y.W.; Investigation: J.L., F.H., Y.L., Y.G., Y.Z, D.C., Z.X.; Methodology: J.L., F.H., D.C.; Project administration: K.X., D.W.; Resources: K.X., Y.W., D.W.; Software: J.L., F.H., Y.G.; Supervision: K.X., Y.W., D.W.; Validation: K.X., Y.W.; Visualization: J.L., F.H., Y.G., K.X.; Writing—original draft: D.W., J.L., F.H.; Writing—review and editing: D.W., J.L., F.H., Y.L., Y.G., Y.Z, D.C., Z.X., Y.W., K.X..

