## Supplementary figures for "Glymphatic System Is Enhanced in Anti-NMDAR Encephalitis and Facilitates Antibody Influx and Clearance in the Brain"

### Supplementary materials

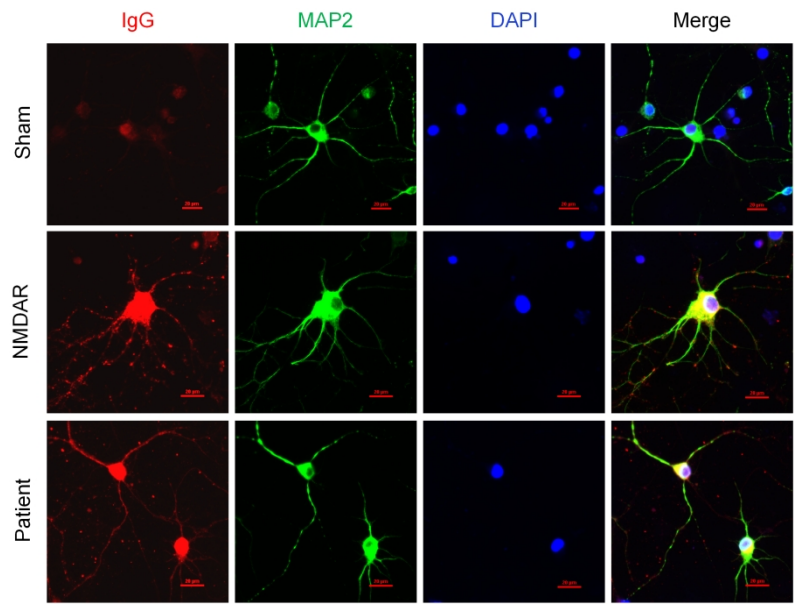

**Figure S1. Serum from anti-NMDAR encephalitis mice shows neuronal binding resembling that of patient-derived anti-NMDAR IgG.** Representative immunofluorescence images of primary rat neurons stained with serum samples from Sham control and GluN1<sub>356-385</sub>-immunized anti-NMDAR encephalitis (NMDAR) mice. Anti-NMDAR IgG prepared from cerebrospinal fluid of the patients was used as a positive control. Bound mouse or human IgG was detected using the corresponding fluorescent secondary antibodies and is shown in red. Green, MAP2; blue, DAPI. Serum from NMDAR mice produced prominent staining with MAP2-positive neuronal somata and neurites, resembling the binding pattern of patient IgG. DAPI, 4',6-Diamidino-2-phenylindole; MAP2, microtubule-associated protein 2. Scale bar, 20 μm.

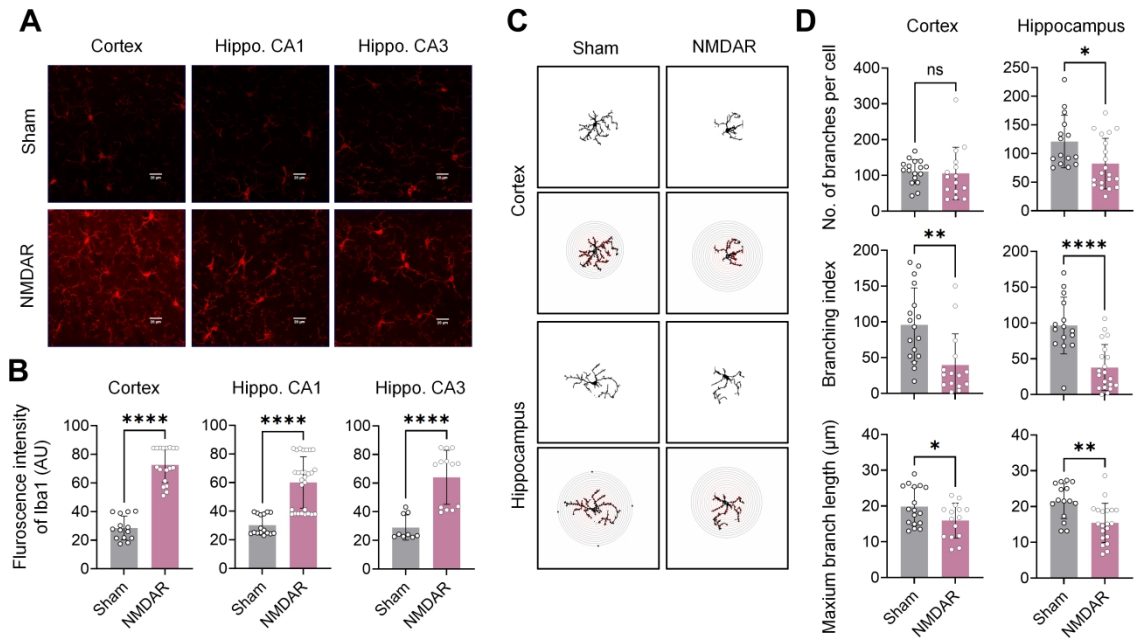

**Figure S2. Activation of microglial cells in anti-NMDAR encephalitis mice.** (A) Immunostaining of Iba1 (red) for microglial cells in the cortex and the CA1 and CA3 regions of the hippocampus from Sham control and GluN1<sub>356-385</sub>-immunized anti-NMDAR encephalitis (NMDAR) mice. Scale bar, 20 μm. (B) Quantification of mean Iba1 fluorescence intensity in the cortex and hippocampal CA1 and CA3 regions (Sham, n = 14, 16, and 9, respectively; NMDAR, n = 18, 28, and 13, respectively). Iba1 fluorescence intensity was higher in all three regions of NMDAR mice than in Sham mice. (C) Representative skeletonized microglial reconstructions and corresponding Sholl analyses from the cortex and hippocampus of Sham and NMDAR mice. (D) Quantification of the number of branches per cell, branching index, and maximum branch length. In the cortex, the number of branches per cell did not differ between groups, whereas the branching index and maximum branch length were lower in NMDAR mice (Sham, n = 17; NMDAR, n = 15). In the hippocampus, all three morphological measures were lower in NMDAR mice than in Sham mice (Sham, n = 15; NMDAR, n = 20). AU, arbitrary unit; Iba1, ionized calcium-binding adapter molecule 1. \*$P < 0.05$, \*\*$P < 0.01$, \*\*\*\*$P < 0.0001$; ns, not significant.

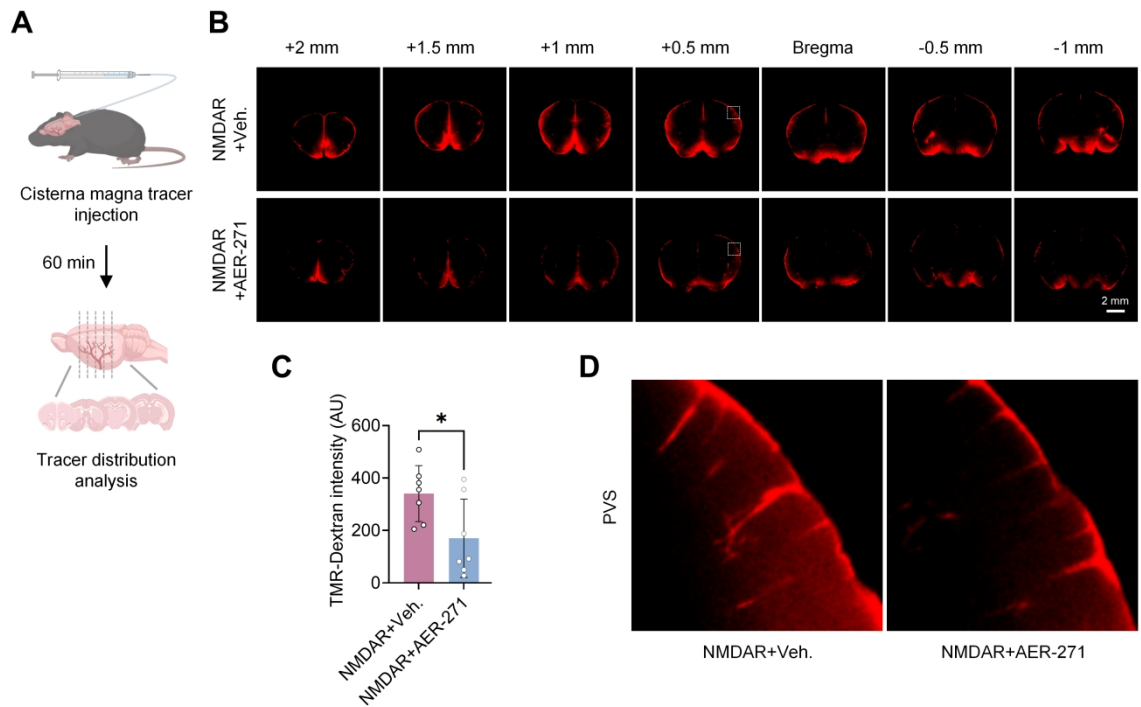

**Figure S3. AER-271 reduces brain influx of TMR-dextran in anti-NMDAR encephalitis mice.** (A) Workflow for ex vivo assessment of tracer influx. TMR-dextran was injected into the cisterna magna. Brains were collected 60 min later, sectioned, and examined for tracer distribution. (B) Representative coronal brain sections showing TMR-dextran (red) distribution from +2.0 to -1.0 mm relative to bregma in vehicle control (Veh.)- and AER-271-treated anti-NMDAR encephalitis mice. Dashed boxes indicate the regions shown at higher magnification in (D). Scale bar, 2 mm. (C) Quantification of TMR-dextran fluorescence intensity across the analyzed coronal sections. The whole-brain section outline was delineated, and the mean fluorescence intensity was calculated after background subtraction, with values from all brain sections of each mouse averaged. Brain TMR-dextran was lower in AER-271-treated mice than in control ($n = 7$). (D) Representative higher-magnification images showing TMR-dextran distribution along cortical perivascular spaces. Perivascular tracer signals were weaker in NMDAR + AER-271 mice than in NMDAR + Veh. mice. Each point represents one mouse. AER-271, 2-(3,5-bis (Trifluoromethyl) phenylcarbamoyl) -4-chlorophenyl dihydrogen phosphate; AU, arbitrary unit; PVS, perivascular spaces; TMR-dextran, tetramethylrhodamine-dextran, 70 kDa. $*P < 0.05$.

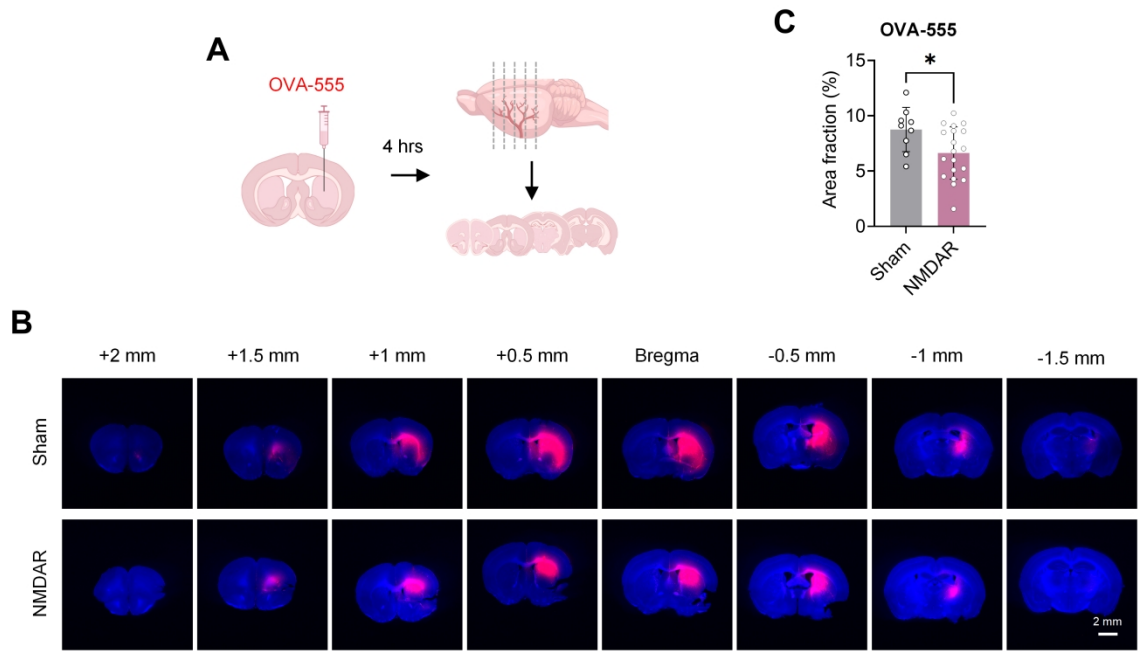

**Figure S4. Brain retention of intrathalamically injected OVA-555 is reduced in anti-NMDAR encephalitis mice.** (A) Workflow for assessing tracer retention in the brain. OVA-555 was stereotactically injected into the thalamus. Brains were collected 4 hours later, sectioned, and examined for residual tracer distribution. (B) Representative coronal brain sections showing OVA-555 distribution from +2.0 to -1.5 mm relative to bregma in Sham and GluN1<sub>356-385</sub>-immunized anti-NMDAR encephalitis (NMDAR) mice. Red, OVA-555; blue, DAPI. Scale bar, 2 mm. (C) Quantification of the OVA-555-positive area fraction across the coronal sections. The ratio of OVA-555-positive area to total brain section area was calculated for each brain section using a unified threshold, and the values from all brain sections of each mouse were averaged. Residual OVA-555 in brain was lower in NMDAR mice than in Sham mice (Sham, $n = 9$; NMDAR, $n = 19$). OVA-555, Alexa Fluor 555 labeled ovalbumin. \*$P < 0.05$.
